# KDM6B inhibition modulates monocyte activation and alleviates IMQ-psoriasis skin inflammation

**DOI:** 10.1101/2025.10.13.682023

**Authors:** Aman Damara, Najla Abassi, Delia Mihoc, Mahsa Nastaranpour, Pauline Kraft, Tina Sarkar, Carsten Deppermann, Johannes U Mayer, Stephan Grabbe, Michael Delacher, Federico Marini, Fatemeh Shahneh

## Abstract

Inflammatory monocytes are increasingly recognized as key amplifiers of psoriasis, yet the epigenetic drivers of their pathogenic signature remain unclear. Here, we demonstrate that the histone demethylase KDM6B is markedly upregulated and catalytically active in classical monocytes during the imiquimod (IMQ)-induced psoriasis model. This is associated with reduced levels of the repressive histone mark H3K27me3, an epigenetic modification linked to chromatin compaction and transcriptional silencing, at the *Il1b, Tnf, Pgam1, Pgk1*, and *Aldoa* promoters, together with an enhanced inflammatory and glycolytic gene signature. Pharmacological blockade of KDM6B after disease onset using GSK-J4, a cell-permeable prodrug that is intracellularly converted to the active KDM6B inhibitor GSK-J1, restores H3K27me3 at inflammatory and metabolic loci, suppresses *Il1b/Tnf* transcription, normalizes bioenergetic profiles, and reduces monocyte and neutrophil recruitment to the inflamed skin.

Single-cell transcriptomic profiling further reveals that KDM6B inhibition represses cytokine-mediated signaling, glycolysis, and chemotaxis pathways in monocytes, yet enriches antigen presentation modules, consistent with a shift toward a homeostatic, antigen-presenting surveillance program in myeloid cells and a Treg-supportive milieu. Collectively, our data identify KDM6B as an epigenetic-metabolic switch that sustains monocyte-driven inflammation in the IMQ-induced psoriasis model. Importantly, we provide preclinical evidence that targeting KDM6B can reduce maladaptive inflammatory response even in progressed diseases. These findings propose KDM6B inhibitors as a promising adjunct to current biologics for psoriasis and other myeloid-driven autoinflammatory disorders.

**Graphical abstract:** 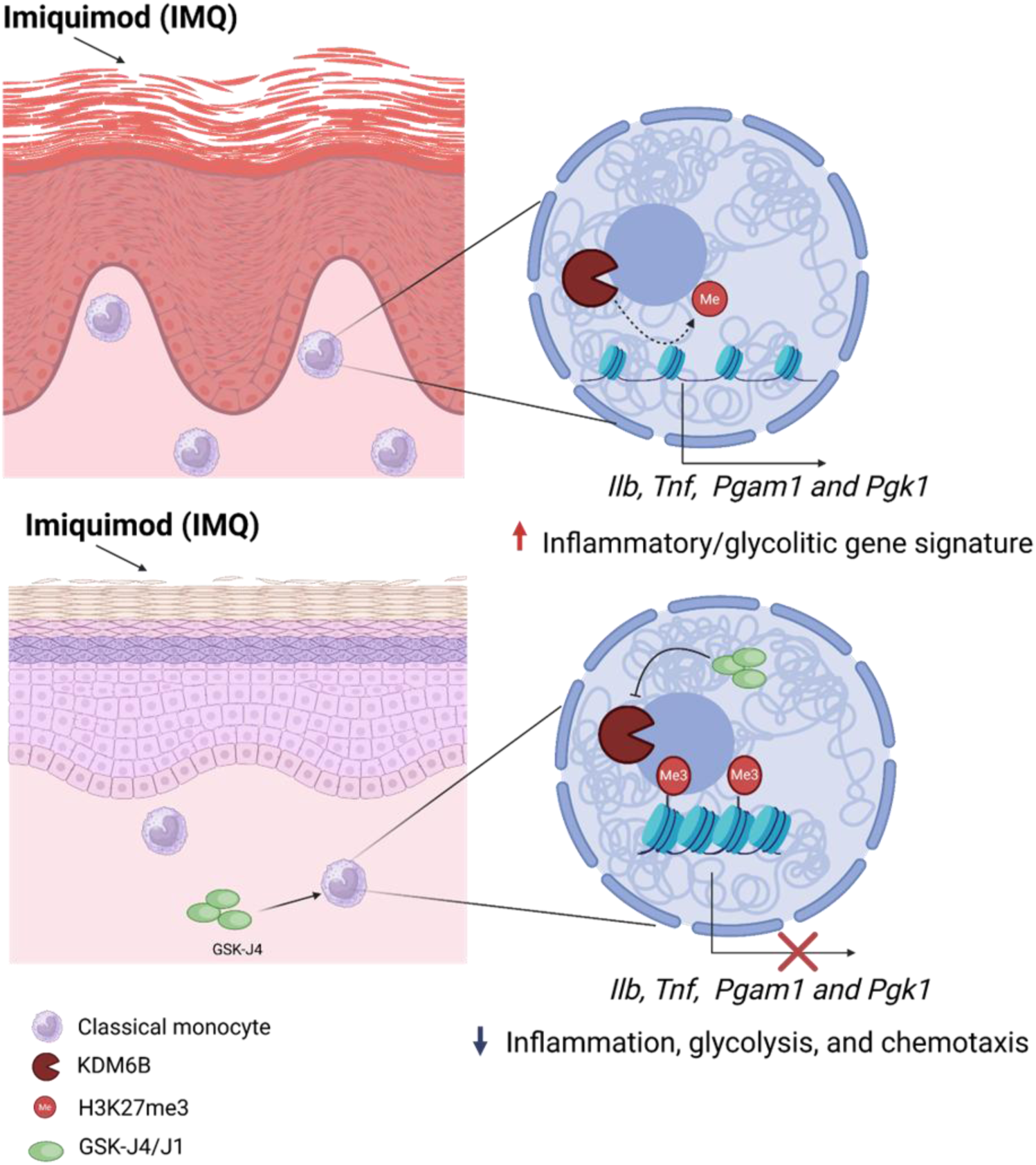

## 1. Introduction

Psoriasis is an autoinflammatory disease characterized by keratinocyte hyperproliferation, aberrant IL-23/IL-17 signaling, and systemic inflammation, affecting 0.1-3% of the global population depending on the region [1,2]. Psoriasis patients are at elevated risk of psoriatic arthritis, cardiometabolic comorbidities, and diminished quality of life [3]. Current biologics that neutralize TNF-α, IL-17A/F, or the IL-23 p19 subunit achieve high short-term response rates, yet up to one third of patients relapse or remain refractory, highlighting the need to define additional disease-sustaining pathways [4].

Emerging evidence highlights the pivotal role of inflammatory monocytes in the pathogenesis of psoriasis [5–7]. In the commonly IMQ-induced psoriasis model, classical monocytes emerge from the bone marrow (BM) and infiltrate psoriatic plaques, where they differentiate into inflammatory macrophages and dendritic cells (DCs) that secrete IL-1β, TNF-α, and IL-6 [5,8–10]. Thus, monocyte recruitment and activation are essential for psoriasis development, and targeting these processes offers promising therapeutic opportunities.

Monocyte phenotypes and function are tightly controlled by transcriptional reprogramming. Epigenetic modifications, such as histone methylation changes and metabolic alterations, including enhanced glycolysis, together determine their inflammatory potential [11,12]. Specifically, the chromatin-modifying enzyme KDM6B, a histone demethylase, removes methyl groups from histone H3K27, thereby reducing the repressive effect of H3K27 trimethylation (H3K27me3) and enabling expression of inflammatory genes in monocytes [13–16]. Elevated KDM6B activity has been shown in monocytes isolated from psoriasis patients [16,17], as well as from individuals with other autoimmune conditions such as rheumatoid arthritis and systemic lupus erythematosus [18,19]. Our recent study also links KDM6B activity to metabolic rewiring toward aerobic glycolysis, further increasing the inflammatory response in isolated monocytes from psoriasis patients [17]. Yet, whether this epigenetic-metabolic switch functions in monocytes in the IMQ-induced psoriasis model and whether its inhibition can reverse established disease remains unresolved. Here, we addressed this knowledge gap with complementary *ex vivo* assays and single-cell RNA-sequencing and examined the KDM6B expression in monocytes using the IMQ-induced psoriasis model [5,8–10]. Mechanistically, monocyte-specific KDM6B promoted NF-κB–mediated transcription of inflammatory genes like *Il1b* and *Tnf* by removing the repressive H3K27me3 from their promoters. Pharmacological inhibition with GSK-J4 attenuated IMQ-induced inflammation and reduced monocyte activation evwn when initiated after disease onset. Single-cell RNA sequencing revealed that KDM6B inhibition suppresses NF-κB–driven pathways and canonical glycolysis in IMQ-treated monocytes. Moreover, we found that IMQ-induced psoriasis is associated with KDM6B-dependent glycolytic reprogramming in classical monocytes, accompanied by loss of H3K27me3 at key glycolytic promoters including *Pgam1*, *Pgk1*, and *Aldoa*. Our findings collectively identified KDM6B as an epigenetic-metabolic checkpoint sustaining psoriatic inflammation and supported KDM6B inhibition as a potential complement to cytokine-targeted therapies in psoriasis and other myeloid-mediated autoinflammatory disorders.

## 2. Methods

### 2.1. Psoriasis-like inflammation model

All mouse experiments were conducted in accordance with the relevant guidelines and regulations for animal welfare issued by the Federal State of Rhineland-Palatinate State Investigation Office (Germany). The study was approved under authorization number G22-1-096, and all efforts were made to minimize animal suffering. Wild-type C57BL/6J mice (both male and female, 20 ± 2 g body weight) were obtained from Jackson Laboratory. Mice were 8-10 weeks old at the start of experiments and housed under specific pathogen-free (SPF) conditions, with controlled temperature and humidity, a 12-hour light/dark cycle, and ad libitum access to food and water. Psoriasis-like skin inflammation was induced using imiquimod (IMQ). Mice received topical applications of 50 mg of 5% IMQ cream (Aldara, Meda, Solna, Sweden) or diluted basis crème (DAC) control (sham) cream daily for five consecutive days on the depilated back skin (3 × 2.5 cm area) and ears. In the IMQ/GSK-J4 group, mice received daily intraperitoneal (*i.p*.) injections of GSK-J4 (1 mg/kg body weight; Med Chem Express, Cat# HY-15648B), starting on day 2 after the first IMQ application. Disease severity was assessed daily using a modified Psoriasis Area and Severity Index (PASI), evaluating three parameters: erythema, scaling, and skin thickness. Each was scored independently on a scale of 0-4 (0 = none, 4 = severe), with a cumulative score ranging from 0 to 12. Body weight was monitored daily from the start of treatment, and weight change was calculated relative to day 1. On day 6, all mice were sacrificed, and samples including spleen, ear, back skin, BM and blood were collected for further analysis. Spleen weights were recorded to assess systemic inflammation. Back skin samples were fixed in 4% formaldehyde (ROTI Histofix, Carl Roth, Germany) for 24 hours, paraffin-embedded, and sectioned (5-6 µm). Sections were stained with hematoxylin and eosin (H&E) and imaged at 10× magnification using an Olympus BX51 microscope.

### 2.2. Peripheral monocytes ablation

Mice were injected *i.p*. every day for up to 6 with 100 mg of anti-mouse CD115-CSF1R (clone: AFS98, Cat# BE0213) and 100 mg anti-mouse Ly6C (clone: Monts 1, Cat# BE0203) to deplete the respective cell populations (classical monocytes from BM) or IgG2a isotype control antibodies (clone: 2A3, Cat# BE0089), respectively. Depleting injections started at day 0, one day before IMQ treatment. The depletion efficacy was tested with staining monocytes against CD45 and CD11b in blood, as shown in **Fig. S1B**.

### 2.3. Cytokine quantification and ELISA

Plasma concentrations of IL-6 (Cat# 431301) were determined using ELISAMAX™ ELISA kits. Assays were performed colorimetrically according to the manufacturer’s instructions. Absorbance was detected at 450 nm using a Hidex Sense microplate reader.

### 2.4. RNA isolation and RT-PCR

On day 6, monocytes were isolated from mouse BM using the EasySep™ Mouse Monocyte Isolation Kit (Cat# 19861A, STEMCELL™ Technologies). Total RNA was isolated from monocytes using the PureLink™ RNA Mini Kit (Cat# 12183025, Thermo Fisher Scientific) according to the manufacturer’s instructions. RNA concentration and purity were quantified using a Nanodrop spectrophotometer (Thermo Fisher Scientific). RNA was reverse-transcribed into cDNA using the High-Capacity cDNA Reverse Transcription Kit (Cat# 4368814, Applied Biosystems). Gene expression was quantified by real-time PCR (CFX384, BioRad) using primers and SYBR Green qPCR Master Mix (Universal) (Cat# HY-K0501A, Med Chem Express). Relative mRNA levels were normalized to the housekeeping gene ACTB and expressed as fold changes using the 2−ΔΔCt method. Primers used for qPCR analysis are as follows: *KDM6B*_F CCTCGTCCTTCCAGGAGTCA, *KDM6B* _R: CTCCTGTAGCTGTGGCTTCC, *Tnf*_F CAGGCGGTGCCTATGTCTCA, *Tnf*_R: GGCTACAGGCTTGTCACTCG, *Il1b* _F: GCCACCTTTTGACAGTGATGAGA, *Il1b* _R: GGACAGCCCAGGTCAAAGGT, *Aldoa* _F 5’-ACA TTG CTG AAG CCC AAC AT-3’, *Aldoa* _R 5’-ACA GGA AAG TGA CCC CAG TG-3*, Eno1*_F 5’-CAT GGG GAA GGG TGT CTC AC-3’, *Eno1*_R 5’-GTG CCG TCC ATC TCG ATC AT-3’, *Gpi1*_F 5’-CAG AGA CAG CAA AGG AGT GG-3’, *Gpi1*_R 5’-GTA GAC AGG GCG ACA AAG TG-3’, *Pgam1*_F 5’-TCT GTG CAG AAG AGA GCA ATC C-3’, *Pgam1*_R 5’-CTG TCA GAC CGC CAT AGT GT-3’, *Pgk1*_F 5’-ATG TCG CTT TCC AAC AAG CTG-3’, *Pgk1*_R 5’-GCT CCA TTG TCC AAG CAG AAT-3’,*Pkm2* _F 5’-TCG CAT GCA GCA CCT GAT T-3’, *and Pkm* _R 5’-CCT CGA ATA GCT GCA AGT GGT A-3’. Gene expression levels were normalized to the *Actin* _F: AGGAGTACGATGAGTCCGGC and *Actin* _R: GGTGTAAAACGCAGCTCAGTA housekeeping gene and presented as fold changes employing the 2−ΔΔCt method.

### 2.5. Chromatin immunoprecipitation (ChIP)

On day 6, monocytes were isolated from mouse BM using the EasySep™ Mouse Monocyte Isolation Kit (Cat# 19861A, STEMCELL™ Technologies). Chromatin immunoprecipitation (ChIP) assays were performed using the Pierce Magnetic ChIP Kit (Cat# 26157, Thermo Fisher Scientific) according to the manufacturer’s protocol. Immunoprecipitation was carried out using 1 µg of anti-H3K27me3 antibody (Cat# C15410195, Diagenode). Normal rabbit IgG served as the negative control for immunoprecipitation. Quantitative PCR (qPCR) analysis was performed using primers targeting promoter regions of the *Il1b* and *Tnf* genes. Primer sequences were as follows: *Il1b*-F 5’-GCA GGA GTG GGT GGG TGA GT-3’; *Il1b*-R 5’-CAG TCT GAT AAT GCC AGG GTG C-3’, *Tnf*-F 5’-TCC TGA TTG GCC CCA GAT TG-3’; *Tnf*-R 5’-TAG TGG CCC TAC ACC TCT GT-3’, *Aldoa* _F 5’-TCC AGG ACA AAT GGG ACT AC-3’, *Aldoa*_R 5’-GGT GGC AGG GCC ACA GCA AA-3’*, Eno1*_F 5’-TAA CGA GCA GGA AAG GAA GAC-3’, *Eno1*_R 5’-AGG AGA GCC TTA AGG ACA GA-3’, *Pgam1*_F 5’-GTG GCA GAG ACA GGA AAT CT-3’, *Pgam1*_R 5’-GAC AGT CTC TCT GCG TAA CC-3’, *Pgk1*_F 5’-GAC AGT CTC TCT GCG TAA CC-3’, *Pgk1*_R 5’-GTG AGA CGT GCT ACT TCC ATT T-3’,*Pkm2* _F 5’-GCA GCC AGC CTG TAA GGG CA-3’, *and Pkm2* _R 5’-GCG AAG ACA GGA AAA CAG TGG GT-3’.

### 2.6. α-Ketoglutarate Quantification

α-Ketoglutarate levels were quantified in monocytes isolated from the BM of four experimental groups of the IMQ-psoriasis mouse model. Measurements were performed using the α-KG Quantitation Kit (Sigma-Aldrich, Cat# MAK541) according to the manufacturer’s instructions.

### 2.7. KDM6B enzymatic activity

Nuclear proteins were extracted from isolated BM monocytes using the Nuclear Extraction Kit (Cat# ab113474, Abcam, Cambridge, UK), following the manufacturer’s protocol. Protein concentrations were subsequently quantified using the Pierce™ BCA Protein Assay Kit (Cat# 23225, Thermo Fisher Scientific, MA, USA). KDM6A/B enzymatic activity was measured using 1 µg of nuclear protein extract per sample with the KDM6A/KDM6B Activity Quantification Assay Kit (Cat# ab156911, Abcam), according to the manufacturer’s instructions.

### 2.8. Seahorse XFp metabolic flux analysis

On day 6, monocytes isolated from BM were seeded at a density of 5 × 10⁵ cells per well in Seahorse XF 8-well mini plates (Cat# 103022-100, Agilent Technologies, CA, USA). Assay medium consisted of XF-RPMI supplemented with 1 mM pyruvate, 2 mM glutamine, and 10 mM glucose, adjusted to pH 7.4. Mitochondrial respiration and glycolytic function were assessed using the XF Cell Mito Stress Test Kit (Cat# 103010-100, Agilent) and the XF Glycolysis Stress Test Kit (Cat# 103020-100, Agilent), respectively. The final working concentrations of inhibitors used for the Cell Mito Stress Test were as follows: 1.5 µM oligomycin, 1 µM FCCP, and 0.5 µM rotenone/antimycin A. For the Glycolysis Stress Test, the following concentrations were used: 10 mM glucose, 1 µM oligomycin, and 50 mM 2-deoxyglucose (2-DG).

### 2.9. Flow Cytometry and Intracellular Staining

On day 6, tissues were collected from mice and processed into single-cell suspensions. Cells were incubated with an unlabeled anti-CD16/32 monoclonal antibody (clone 93, Cat# 14-0161-82, Invitrogen) for 10 minutes at 4°C to block non-specific Fc receptor binding. After washing with FACS buffer (PBS containing 2% FBS and 2 mM EDTA), the cell pellets were resuspended in a fixable viability dye along with fluorochrome-conjugated surface antibodies and incubated for 30 minutes at 4°C.

Monocytes were identified by surface staining with the following antibodies: CD11b (Cat# 101212, clone M1/70, APC, BioLegend), CD45 (Cat# 103147, clone 30-F11, BV711, BioLegend), Ly-6G (Cat# 127618, clone 1A8, PE-Cy7, BioLegend), and Ly-6C (Cat# 128012, clone HK1.4, PerCP-Cy5.5, BioLegend). To exclude T and B cells (Lin) from the analysis, anti-CD3 (Cat# 100233, clone 17A2, BV510, BioLegend) and anti-CD45R (Cat# 103248, clone RA3-6B2, BV510, BioLegend) antibodies were included. For T cell analysis, cells were stained with antibodies against CD4 (Cat# 100443, clone GK1.5, BV421, BioLegend), CD8a (Cat# 100714, clone 53-6.7, APC-Cy7, BioLegend), CD45 (Cat# 103147, clone 30-F11, BV711,

BioLegend), and TCRγδ (Cat# 18123, clone GL3, PE-Cy7, BioLegend). Following surface staining, cells were fixed and permeabilized using the Foxp3/Transcription Factor Staining Buffer Set (Invitrogen, Cat# 5523), and intracellular staining was performed using fluorochrome-conjugated antibodies. For monocytes, intracellular markers included anti-TNFα (Cat# 506328, clone MP6-XT22, BV421, BioLegend), anti-Pro-IL1β (Cat# 12711482, clone NJTEN3, PE, Invitrogen), and anti-KDM6B (Cat# NBP1-06640AF488, AF488, Bio-Techne GmbH). T cells were intracellularly stained with antibodies against RORγT (Cat# 562894, clone Q31-378, BV421, BioLegend), and Foxp3 (Cat# 12-5773-82, clone FJK-16s, PE, eBioscience). All samples were acquired using a BD LSRII flow cytometer and analyzed using FlowJo v10.10.0 software (BD Biosciences).

### 2.10. Single-cell RNA Sequencing

C57BL/6 female mice were randomly assigned in duplicates to three experimental groups: Sham, IMQ, and IMQ/GSK-J4, and treated as described above. On day 6, spleen and skin tissues were harvested and further processed to generate single cell suspensions. Spleen was disrupted onto a filter mesh in a 3 cm Petri dish. Red blood cells were lysed using commercial RBC lysis buffer by incubation at room temperature for 3 minutes, followed by centrifugation for 2 minutes at 4°C and 1000 rcf. Dorsal skin was excised, manually cut into small pieces using scissors and digested in DMEM (Gibco #11960044), 4 mg/ml collagenase type IV (Sigma-Aldrich #C5138), 20 μg/ml DNase I (Thermo Fisher Scientific #89836) and 5 mg/ml bovine serum albumin (Carl Roth #90604-29-8). Skin tissue was digested using the gentleMACS C tubes (Miltenyi Biotec) and the program ‘37C_Multi_H’’ on the gentleMACS dissociator (Miltenyi Biotec) was performed. Murine samples were stained using the following surface anti-mouse antibodies: CD45-BUV737 (BD Biosciences #752414), CD4-BV421 (Biolegend #566644), CD8-PE-Cy7 (Biolegend #100722), CD25-PE (Biolegend #102008), CD11b-APC (Biolegend #101212), CD11c-APC (Biolegend #117310), CD19-APC (Biolegend #115512), MHCII-APC (Biolegend #107614), CD206-APC (Biolegend #141708), NK1.1-APC (Biolegend #108710) and Zombie NIR (Biolegend #423106). Individual samples were stained with 6 TotalSeq™-C anti-mouse hashtag antibodies (Biolegend C1 #155861, C2 #155863, C3 #155865, C4 #155867, C5 #155869, C6 #155871) at 4°C for 20 minutes in 100 μL FACS buffer (StemCell Technologies #07905). CD45⁺ immune cells were sorted for single-cell RNA sequencing. The following numbers of cells were sorted into 1.5 mL Eppendorf tubes containing 1× PBS supplemented with 0.5% BSA and maintained at 4°C: 8,000 CD45⁺ cells from spleen (Sham: TotalSeqC1; IMQ: TotalSeqC3; IMQ/GSK-J4: TotalSeqC5), and 4,000 CD45⁺ cells from skin (Sham: TotalSeqC2; IMQ: TotalSeqC4; IMQ/GSK-J4: TotalSeqC6). Following sorting, cells were centrifuged at 300 × g for 5 minutes at 4°C, and the supernatant was carefully removed. The resulting cell pellets were resuspended in 37.5 µL of 0.5% BSA-PBS buffer. Cell suspensions were mixed with Master Mix from the Chromium Next GEM Single Cell 5′ v2 (Dual Index) kit with Feature Barcode Technology (10x Genomics, Cat# 1000266), and samples were loaded onto a Chromium Next GEM Chip K (10x Genomics, Cat# 2000182). Single-cell libraries were prepared according to the manufacturer’s protocol (Chromium Next GEM Single Cell 5′ v2 with Feature Barcode Technology, CG000330 Rev F). Prepared scRNA libraries were sequenced using an Illumina NextSeq™ 550 system with a 150-cycle high-output cartridge.

### 2.11. Bioinformatics Analysis

Initial data processing, quality control, and integration followed the previously established protocol described by Nedwed et al [20]. Raw sequencing reads were demultiplexed and aligned using Cell Ranger v7.1.0 (10x Genomics) [21], employing a transcriptome index generated from the Mus musculus genome build GRCm38. Gene annotation was based on ENSEMBL release 102 for Mus musculus. Quality control (QC) was performed independently on each dataset to remove poor-quality cells using scater v1.32.0 [22]. Mitochondrial gene content served as a proxy for identifying damaged cells, using a threshold of three median absolute deviations above the median, in accordance with recommendations from the OSCA guidelines [23]. Doublet detection was performed with scDblFinder v1.18.0 [24]. After QC, normalization of cell-specific biases was conducted using the deconvolution-based method implemented in scran [25]. Counts were divided by size factors, and normalized values were log-transformed after adding a pseudocount of one. Integration of datasets from different biological samples was achieved using the Mutual Nearest Neighbors (MNN) method implemented in batchelor v1.20.0 [25]. Subsequently, highly variable genes (HVGs) were nidentified by decomposing per-gene variability into technical and biological components based on the mean-variance trend. Dimensionality reduction was performed using Principal Component Analysis (PCA). PCA results were then used as input for t-distributed stochastic neighbor embedding (t-SNE) for visualization [26]. Clustering analysis utilized the 9,982 most highly variable genes to build a shared nearest neighbor graph (SNNG) [27]. Clusters were determined using the Louvain community detection algorithm implemented via igraph (https://doi.org/10.5281/zenodo.7682609).

Initial automated cell-type annotation was performed using SingleR v2.6.0 [28] with the ImmGenData reference from celldex v1.14.0 [28]. These annotations were manually refined using established cell-type marker genes from the literature. Visualization and manual refinement of cell clusters were carried out using iSEE v2.16.0 [29] and iSEEfier v1.2.0 (https://bioconductor.org/packages/iSEEfier). Most visualizations for single-cell data were generated using iSEE. Pseudobulk differential gene expression analysis was conducted using muscat v1.20.0 [30] with the default modeling approach. Multiple testing corrections were applied using the Benjamini-Hochberg (BH) method, and genes with adjusted p-values < 0.05 were considered DEGs. Functional enrichment analysis of DEGs was performed with mosdef v1.2.0 (https://bioconductor.org/packages/mosdef), implementing the topGO package [31]. Enrichment analysis used the “elim” algorithm within the Biological Process ontology, employing DEGs as input against a background set comprising all detected genes. Finally, differential abundance analysis comparing cell type proportions across treatment conditions was performed using speckle v1.6.0 [32]. Statistical significance was defined at a False Discovery Rate (FDR) < 0.05.

### 2.12. Statistical Analysis

The sample size, number of replicates, and detailed statistical information for each experiment are provided in the figure legends. All data are presented as mean ± SEM, as specified in the respective figure legends unless otherwise stated. Shapiro–Wilk test was used to determine normality of data, Brown-Forsythe or F test to determine variances. Parametric statistical analysis was performed using one-way ANOVA followed by post hoc analysis (Newman–Keuls multiple comparison test) for analysis of pairwise differences between more than groups. Significance was evaluated by the above-mentioned tests and was performed in GraphPad Prism (v6).

## 3. Results

### 3.1. Monocyte depletion reduces IMQ-induced psoriasis severity

To better understand the role of monocytes in the IMQ-induced psoriasis model, we first examined how depleting monocytes influences disease onset and severity. Mice were treated daily with an intraperitoneal injection of depleting antibodies directed against CSF1R (CD115) and Ly6C (100 µg each per mouse) or an equivalent dose of isotype IgG starting one day before topical challenge (day 0) throughout the experiment **(Fig. 1A and S1A)**. From day 1 to day 5, 5 % IMQ cream (Aldara) or the vehicle control DAC cream was applied once daily to both ears (10 mg per ear) and the shaved dorsal skin (50 mg) **(Fig. 1A)**. Monocyte depletion in the periphery markedly ameliorated skin inflammation on day 6 with reduced erythema and scaling **(Fig. 1B-C)**, and significantly decreased spleen size **(Fig. 1D)**, consistent with attenuated systemic immune activation. Histological analysis revealed a significant reduction in epidermal thickness and inflammatory cell infiltration in IMQ/monocyte-depleted mice compared to IMQ alone **(Fig. 1B)**. Clinical scoring over six days revealed significantly reduced erythema, scaling, and skin thickening in the monocyte-depleted group **(Fig. 1B-C)**. Furthermore, plasma cytokine analysis by ELISA showed significantly lower levels of IL-6, a key pro-inflammatory cytokine elevated in the IMQ model, in monocyte-depleted mice compared to IMQ controls **(Fig. 1E)**. Flow cytometry demonstrated decreased frequencies of CD11b⁺Ly6G⁻ monocytes in blood and skin **(Fig. 1F)**, supporting the notion that monocytes are critical drivers not only of local skin pathology but also of systemic inflammatory responses.

**Fig. 1.**
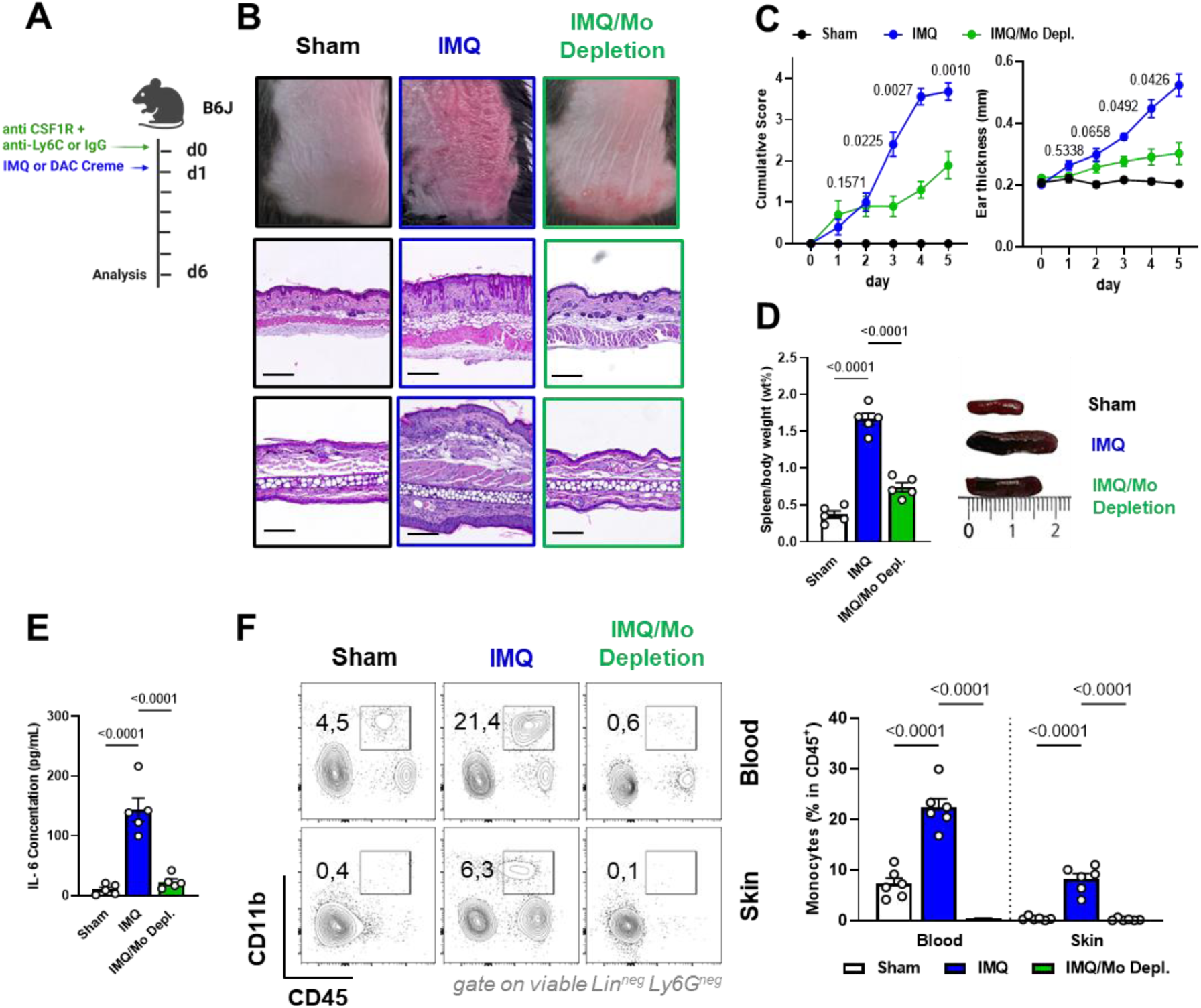
Classical monocytes drive inflammation in psoriatic skin and systemic compartments in IMQ-psoriasis mouse model. (A) Experimental schematic for *in vivo* monocyte depletion via anti-CSF1R and anti-Ly6C antibodies followed by IMQ application. (B) Representative images and H&E-stained skin sections from IMQ-treated and IMQ/monocyte-depleted mice. Scale bars: 100–200 µm. (C) Cumulative score assessing erythema and scaling (left) as well as quantification of ear thickness (right) over six days of IMQ treatment. (D) Spleen weight was significantly reduced upon monocyte depletion. (E) ELISA analysis of plasma IL-6 levels in monocyte-depleted animals compared to IMQ-treated and sham-treated mice. (F) Representative flow cytometry plots showing frequencies of CD11b⁺Ly6G⁻ monocytes in blood and skin. Data are representative of 2 independent experiments (n = 6). Data are presented as mean ± SEM; statistical significance was assessed using one-way ANOVA with Newman–Keuls multiple comparison test.

### 3.2. KDM6B expression is elevated in monocytes from the IMQ-induced psoriasis model

To dissect the cellular landscape and functional states of immune cells in the IMQ-induced psoriasis model, we performed single-cell RNA sequencing (scRNA-seq) on CD45^+^ cells isolated from the skin and spleen of IMQ-treated and sham-treated mice. Dimensionality reduction using t-distributed stochastic neighbor embedding (t-SNE) revealed transcriptionally distinct clusters corresponding to major immune cell types, including classical (*Itgam*, *Ly6c2*, *Ly6g^neg^*) and non-classical monocytes (*Itgam*, *Spn*, *Ly6c2^low^*), neutrophils (*Ly6g*), macrophages (*Adgre1*, *Fcgr1*), dendritic cells (for whole DCs: *Zbtb46 and Itgax*, cDC1: *Xcr1*; cDC2: *Sirpa,* moDCs: *Cd209*, *H2-Ab1*), CD4⁺ and CD8⁺ T cells (*Cd3e, Cd4, Cd8a*), Tregs (*Foxp3*), Th17 cells (*Il17a, Rorc*), NK cells (*Ncr1*), ILCs (*Gata3, Rorc*), and B cells (*Cd19*, *Cd20*) **(Fig. 2A and S2B)**.

**Fig. 2.**
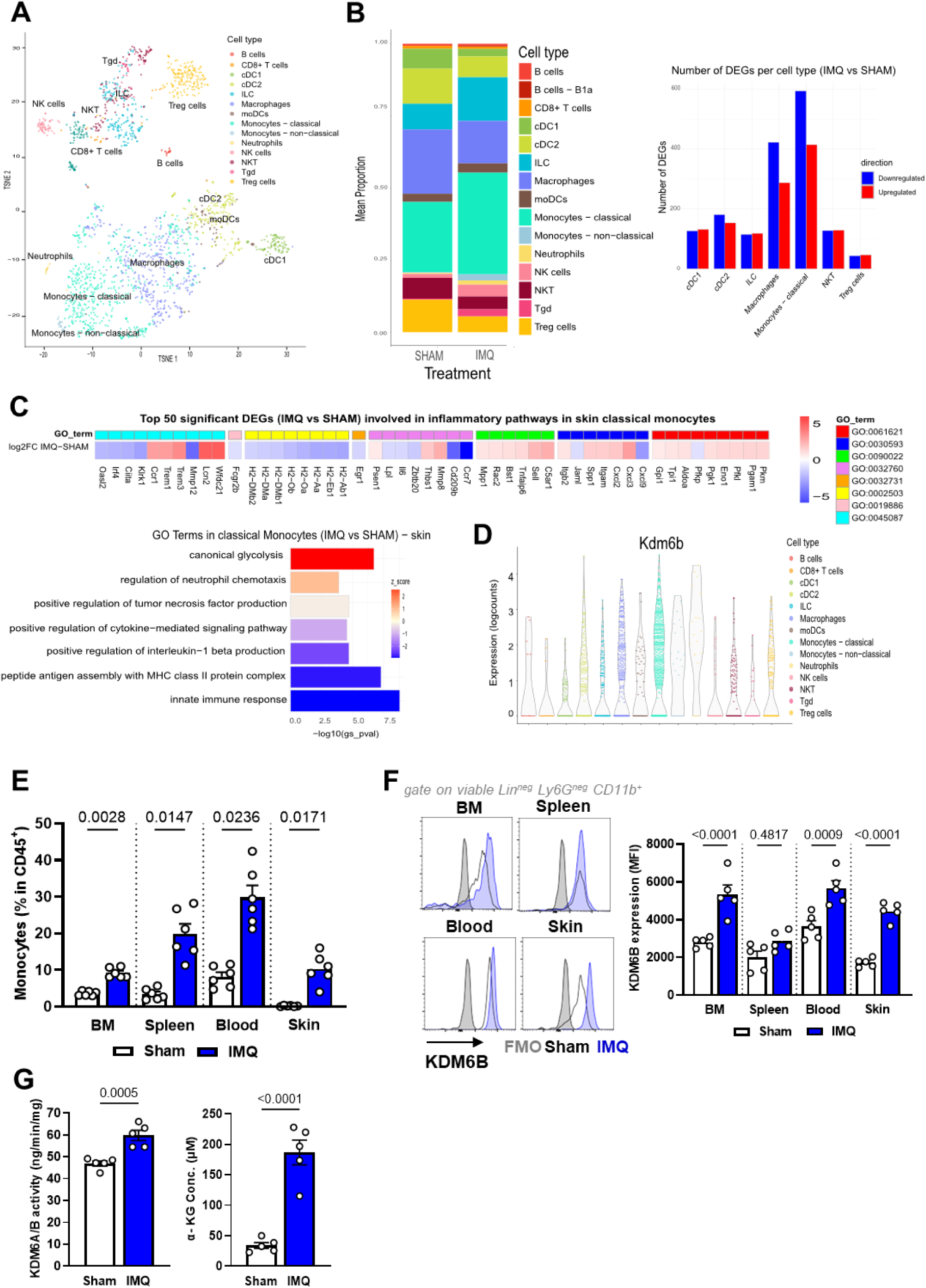
KDM6B expression and α-KG levels are increased in monocytes in the IMQ-psoriasis mouse model. (A) t-SNE plot of CD45⁺ immune cells isolated from the skin of IMQ-treated and sham mice. (B) Relative abundance of immune cell subsets in sham-treated versus IMQ-treated skin (left) and number of differentially expressed genes (DEGs) across immune populations (right). (C) Differential gene expression and GO enrichment analysis of DEGs in classical monocytes from IMQ versus sham skin. (D) KDM6B expression in immune cell clusters from skin of IMQ-treated versus sham-treated mice. (E) Representative flow cytometry plots showing frequencies of CD11b⁺Ly6G⁻ monocytes in BM, spleen, blood, and skin. (F) Quantification of KDM6B protein expression by flow cytometry. Data are presented as normalized modal mean fluorescence intensity (MFI). (G) Quantification of intracellular KDM6B activity and α-KG levels in monocytes from IMQ-treated versus sham-treated mice by ELISA. Data are representative of three independent experiments (n = 5-6). Statistical analysis was performed using an unpaired t-test. Data are presented as the mean ± SEM.

Comparison of immune cell frequencies between IMQ-treated and sham-treated mice demonstrated that classical monocytes were markedly increased in the skin of IMQ-treated mice compared to the spleen **(Fig. 2B and S2C)**. Furthermore, differential expression analysis revealed substantial transcriptional changes in classical monocytes and macrophages, with both upregulated and downregulated genes, indicating a strong myeloid bias in the inflamed skin **(Fig. 2B)**. Differential gene expression analysis of classical monocytes from IMQ-treated versus sham-treated skin revealed upregulation of genes involved in inflammatory and metabolic pathways **(Fig. 2C)**. Gene Ontology (GO) enrichment demonstrated significant activation of canonical glycolysis (GO:0061621), neutrophil chemotaxis (GO:0030593 and GO:0090022), TNF production (GO:0032760), and IL-1β signaling (GO:0032731). Notably, despite the overall inflammatory profile, pathways related to antigen processing via MHC class II (GO:0002503 and GO:0019886) and innate immune responses (GO:0045087) were significantly downregulated in classical monocytes from IMQ-treated skin, suggesting a shift away from antigen presentation toward a metabolically and cytokine-driven inflammatory phenotype **(Fig. 2C)**.

Based on our previous observations in peripheral monocytes from psoriatic patients showing KDM6B upregulation [16,17], we next examined KDM6B expression in the IMQ-induced psoriasis model. Consistent with the human data, KDM6B expression was elevated in the classical monocyte and myeloid clusters in IMQ-treated skin compared to sham-treated skin and the other clusters **(Fig. 2D)**. To validate these findings systematically, we analyzed monocyte frequencies and KDM6B expression across multiple tissues [5,8,10]. Monocytes were isolated from BM, spleen, blood, and skin of mice treated with IMQ for six days, as well as sham controls, for subsequent analyses. Flow cytometry revealed a significant increase in CD11b⁺Ly6G⁻ monocyte frequencies among CD45⁺ cells **(Fig. 2E and S2D)**. Importantly, KDM6B protein expression was elevated in CD11b⁺ monocytes (gated on viable CD45⁺Lin⁻Ly6G⁻ cells) across all tissues analyzed compared to controls **(Fig. 2F)**. Additionally, we observed significantly increased KDM6B enzymatic activity together with elevated intracellular α-ketoglutarate (α-KG) levels, a key metabolic cofactor required for KDM6B demethylase function [33], in BM monocytes **(Fig. 2G)**. These findings support the notion that KDM6B is upregulated in monocytes during psoriasis-like inflammation, coupled to increased availability of its metabolic cofactor α-KG.

### 3.3. Systemic inflammation is reduced by GSK-J4 treatment in a pre-established IMQ-psoriasis mouse model

Small-molecule inhibitors specifically targeting KDM6B, such as GSK-J4 [34,35], have recently been developed for inflammatory diseases like SLE and abdominal aortic aneurysms, showing promising efficacy both *in vitro* and *in vivo* [14,18]. However, it remains unclear whether KDM6B plays a critical role in psoriasis-induced inflammation in the IMQ-induced psoriasis model and whether KDM6B inhibition could effectively alleviate an already established skin inflammation. To address this question, mice were treated with IMQ-containing cream for five consecutive days. Beginning on day 2 of the IMQ course (after clinical onset), mice additionally received intraperitoneal injections of either GSK-J4 (10 mg/kg) or vehicle control (DMSO) **(Fig. 3A)**. Systemic administration of GSK-J4 significantly reduced ear thickness and overall psoriasis severity, reflected by lower composite clinical scores (erythema, scaling, thickness; modified PASI) **(Fig. 3B and 3C)**. Histological analysis of the back skin and ears revealed that GSK-J4 treatment markedly reduced epidermal hyperplasia and leukocyte infiltration, preserving epidermal architecture **(Fig. 3C)**. Moreover, plasma IL-6, a systemic pro-inflammatory mediator implicated in keratinocyte proliferation [1], was decreased **(Fig. 3D)**. GSK-J4 treatment also attenuated splenomegaly in IMQ-treated mice **(Fig. 3E)**. Collectively, these findings indicate that inhibiting KDM6B effectively attenuates both systemic and local skin inflammation induced by IMQ in mice.

**Fig. 3.**
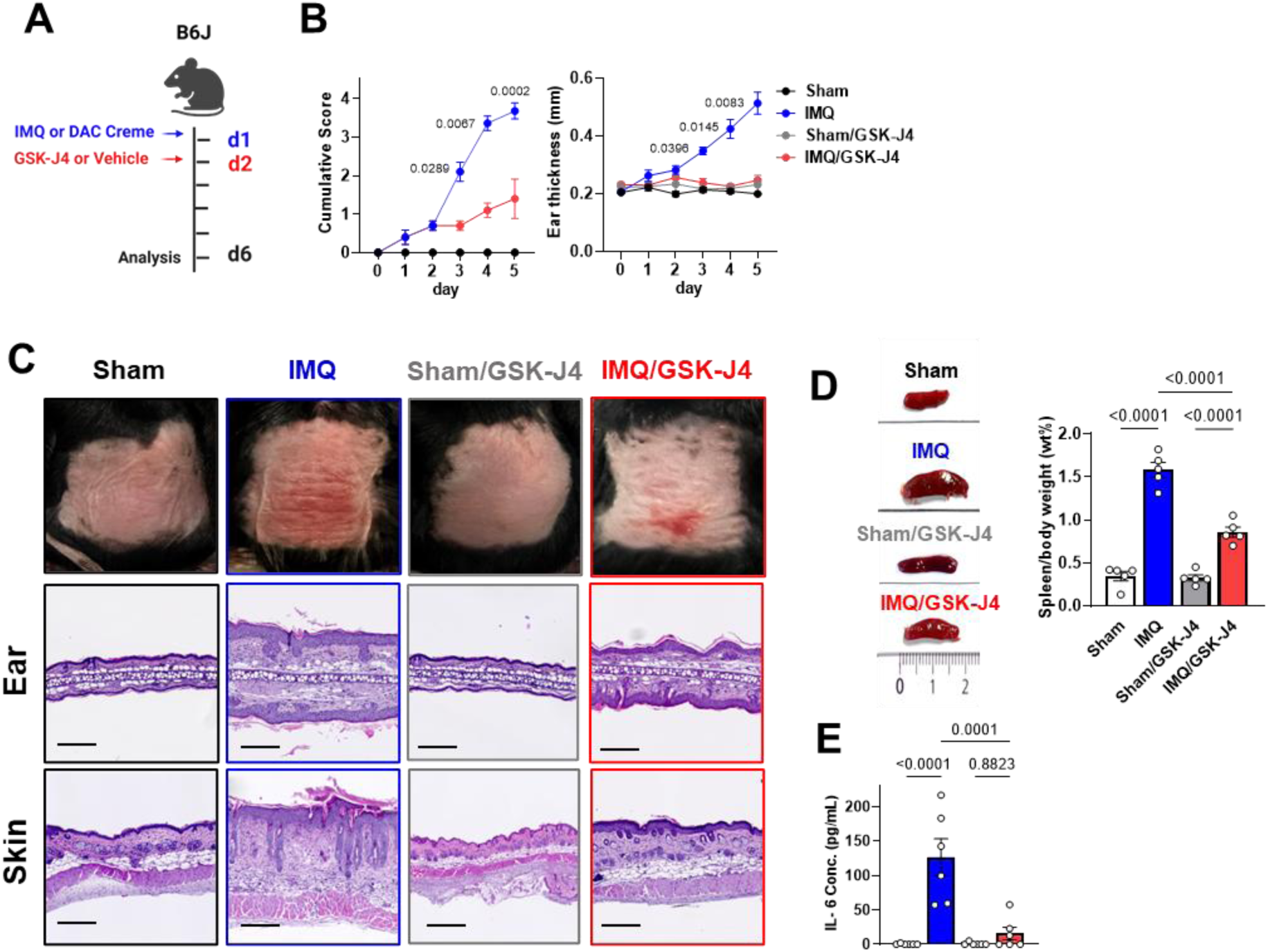
GSK-J4 treatment decreases inflammation in psoriasis severity and infiltration of inflammatory monocytes into the skin. (A) Treatment scheme. Intraperitoneal vehicle or GSK-J4 injection started on day 2 of the IMQ course (post-onset). (B) Quantification of ear thickness, erythema, and scaling (PASI score) throughout the experiment (n = 12). (C) Representative macroscopic images of sham-treated, IMQ-treated, and IMQ-induced psoriasis treated with GSK-J4. H&E staining of ears and dorsal skin on day 6. Scale bars: 100 µm. (D) Quantification of the spleen-to-body weight ratio across treatment groups. (E) Plasma IL-6 levels from all treatment groups were quantified on day 6 using ELISA (n = 6-8). Data are representative of 4 independent experiments. Data are presented as mean ± SEM; statistical significance was assessed using one-way ANOVA with the Newman-Keuls multiple comparison test.

### 3.4. Myeloid cell infiltration is reduced by GSK-J4 treatment in the IMQ-psoriasis mouse model

Myeloid cell infiltration into the skin and peripheral tissues is a hallmark of IMQ-psoriasis and is closely associated with disease progression [5,6]. To understand how inhibiting KDM6B ameliorates psoriasis severity, we examined the impact of pharmacological KDM6B inhibition using GSK-J4 on immune cell infiltration and inflammatory cytokines expressed by monocytes in IMQ-treated mice. Flow cytometry analysis revealed that IMQ-induced psoriasis increased the abundance of neutrophils (viable CD45⁺Lin⁻ [CD3^−^CD19^−^Nk1.1^−^]CD11b⁺Ly6G⁺) **(Fig. 4A)** and classical monocytes (viable CD45⁺Lin⁻Ly6G⁻CD11b⁺Ly6C⁺) **(Fig. 4B)** in both blood and skin, which were significantly reduced following GSK-J4 treatment. These findings indicate that KDM6B inhibition limits pathogenic myeloid infiltration during active IMQ-induced psoriatic inflammation. In addition, intracellular staining of monocytes showed notably lower levels of TNF-α and IL-1β proteins in both blood and skin of IMQ/GSK-J4-treated mice compared with IMQ-treated mice **(Fig. 4C)**. These reductions in key proinflammatory cytokines suggest that GSK-J4 not only limits myeloid infiltration but also dampens their inflammatory potential.

**Fig. 4.**
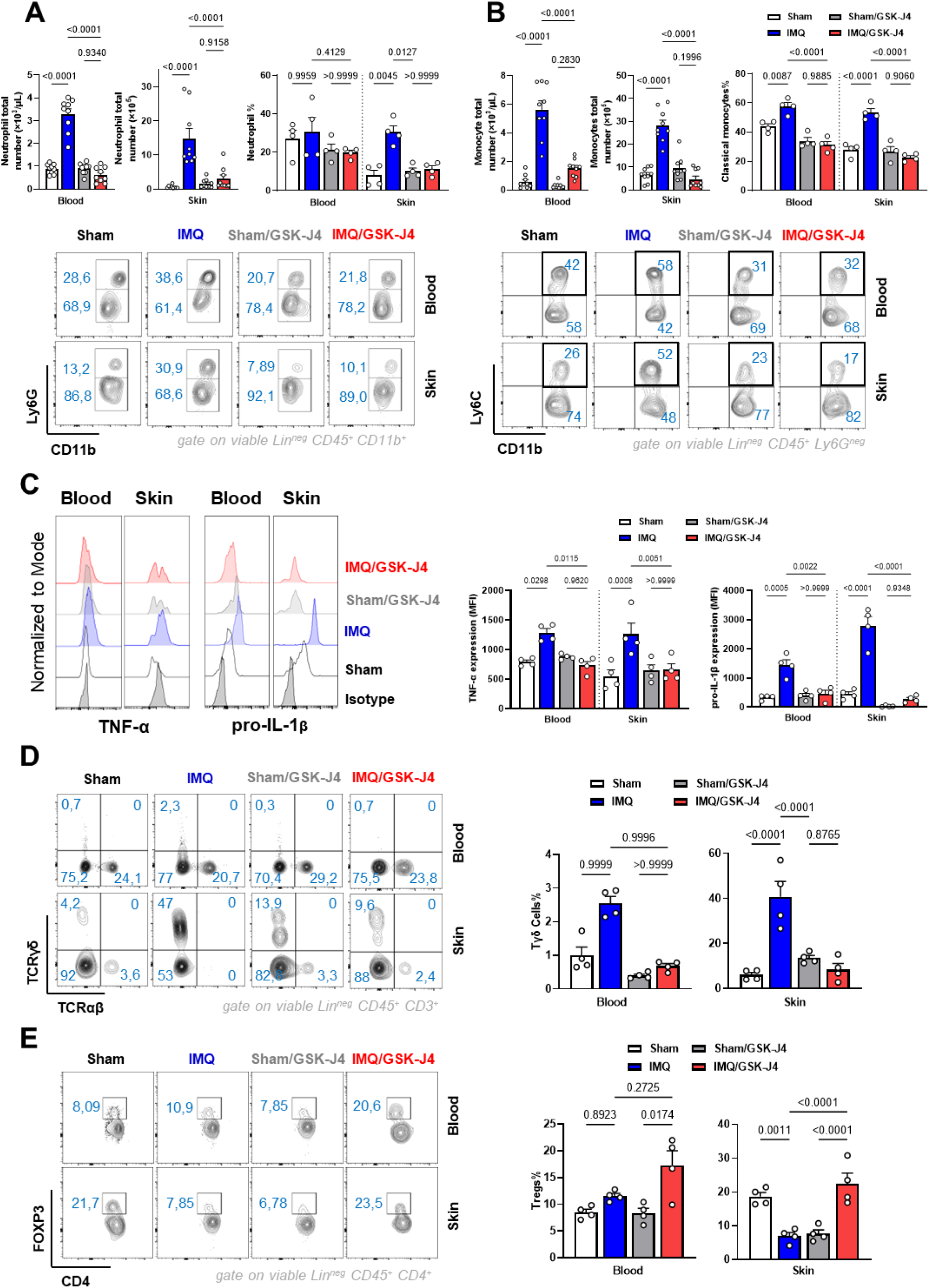
GSK-J4 treatment decreases infiltration of inflammatory monocytes into the skin in established IMQ-psoriasis. (A) Total numbers and percentages of neutrophils in the blood and skin of sham-, IMQ-, sham/GSK-J4-, and IMQ/GSK-J4-treated mice, quantified by flow cytometry (n = 6–8). (B) Total numbers and percentages of classical monocytes in the same groups, quantified by flow cytometry (n = 6–8). (C) Intracellular expression of TNF-α and IL-1β in CD11b⁺Ly6G⁻ monocytes were measured in blood and skin after *ex vivo* stimulation. Data are presented as normalized modal mean fluorescence intensity (MFI). (D) Frequencies of γδ T cells in blood and skin were assessed by intracellular cytokine staining. (E) Flow cytometric quantification of CD4⁺FOXP3⁺ Tregs in blood and skin. Data are representative of four independent experiments (n = 4–6) and are presented as mean ± SEM. Statistical significance was determined using one-way ANOVA with the Newman-Keuls multiple comparison test.

Since monocytes have been shown to shape downstream adaptive immune responses, including γδ T cells and regulatory T cells (Tregs) in the IMQ-psoriasis mouse model [5,36], we examined these populations and observed a significant decrease in γδ T cells (gated on viable CD45⁺CD3⁺TCRαβ⁻TCRγδ⁺ cells) in the skin following GSK-J4 treatment **(Fig. 4D)**. This reduction may reflect an indirect consequence of suppressed IL-1β production by myeloid cells, as IL-1β is a potent inducer of γδ T cells [5]. Importantly, GSK-J4 treatment led to a significant increase in regulatory CD4⁺FOXP3⁺ Tregs (gated as viable CD45⁺CD4⁺FOXP3⁺) in the blood and skin **(Fig. 4E)**, suggesting a shift toward an immunoregulatory milieu. This observation supports the hypothesis that epigenetic modulation via KDM6B inhibition not only reduces inflammatory responses but also promotes immune regulation [37–39]. However, the possibility that GSK-J4 directly influences γδ T cells and Tregs cannot be excluded. Together, these results suggest that GSK-J4 attenuates psoriasis-like inflammation by suppressing myeloid cell infiltration, proinflammatory cytokine production, and the frequency of γδ T cells, while enhancing Tregs numbers. While our data identifies monocytes as the primary cellular targets of GSK-J4 treatment, KDM6B inhibition broadly reshapes the inflammatory cell components in psoriatic-like skin and contributes to a broad anti-inflammatory response. These findings highlight the therapeutic potential of targeting KDM6B in psoriasis and other inflammation-driven diseases characterized by aberrant myeloid activation.

### 3.5. KDM6B inhibition by GSK-J4 causes remodeling of classical monocyte transcriptional programs in the IMQ-psoriasis mouse model

Recent studies have demonstrated that GSK-J4 significantly reduces the expression of proinflammatory cytokine genes, including *Il6*, *Il1b*, *Tnf*, and *Il12* in monocytes and macrophages [14,18,39–41]. To comprehensively understand the immune landscape and the broader impact of systemic KDM6B inhibition on immune cells in IMQ-treated mice, we performed scRNA-seq on isolated CD45^+^ immune cells from the skin and spleen of sham-treated, IMQ-treated, and IMQ/GSK-J4-treated mice. Dimensionality reduction using t-SNE revealed major immune cell subsets, including classical and non-classical monocytes, neutrophils, macrophages, DCs, CD4⁺ and CD8⁺ T cells, Tregs, NK cells, and B cells in the skin **(Fig. 5A)** and spleen **(Fig. S2A)**. Notably, GSK-J4 treatment led to a significant reduction in classical monocyte abundance in the skin, partially restoring the cellular composition toward the homeostatic distribution observed in sham-treated skin **(Fig. 5A)**.

**Fig. 5.**
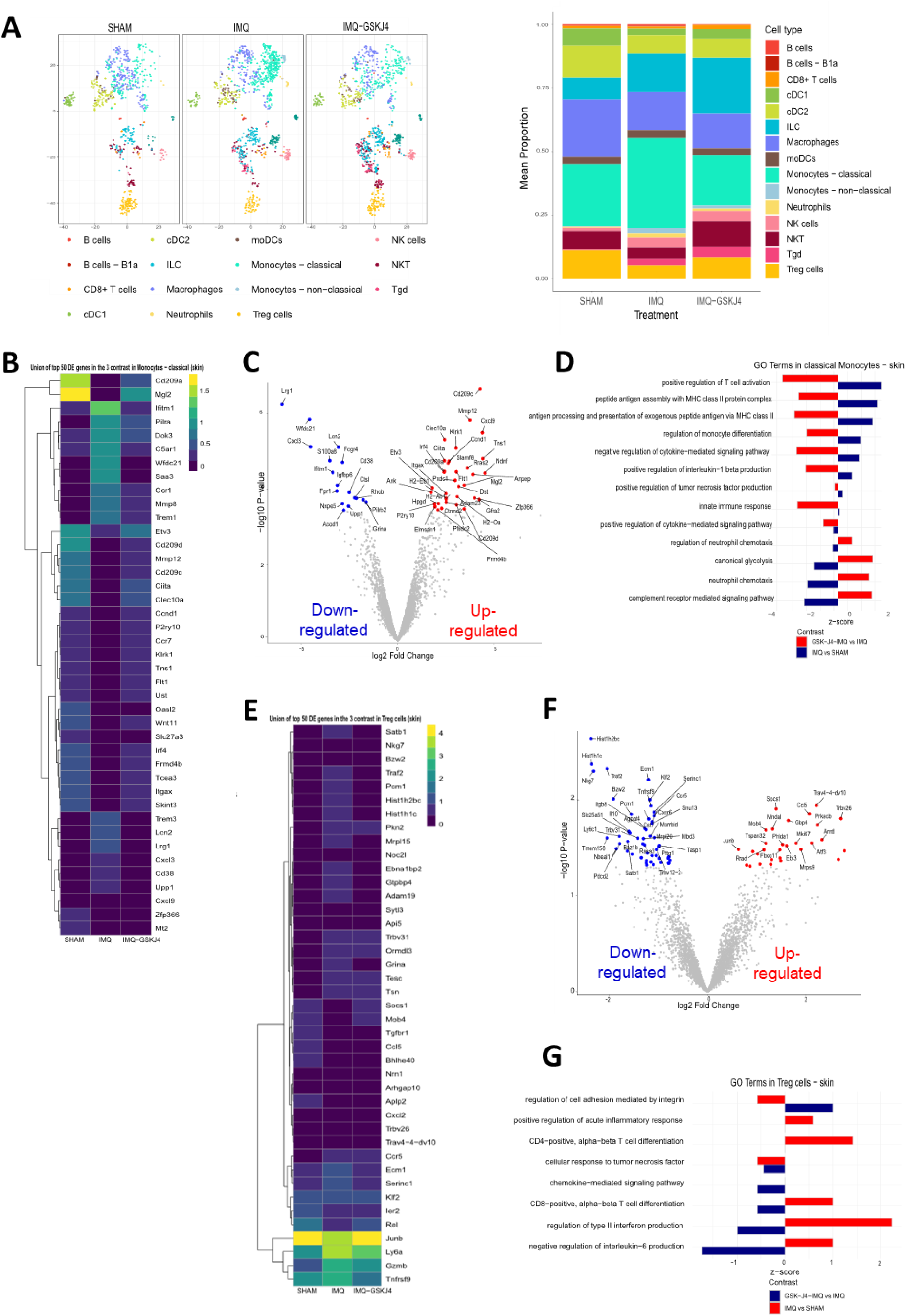
Single-cell transcriptional profiling reveals reduced inflammatory pathway expression in classical monocytes following GSK-J4 treatment in IMQ-psoriasis mouse model. (A) t-SNE plots showing major immune cell populations identified from CD45⁺ cells isolated from the skin of sham-, IMQ-, and IMQ/GSK-J4-treated mice. Each dot represents a single cell, color-coded by experimental condition. Stacked bar plot showing the relative abundance of immune cell subsets. (B) Heatmap showing pseudobulk expression of the top 50 DEGs in classical monocytes across sham, IMQ, and IMQ/GSK-J4 groups; filtered heatmap of adjusted-P DEGs (n = 53) shown as indicated. (C) Volcano plot of DEGs in classical monocytes comparing IMQ/GSK-J4 versus IMQ groups. (D) GO enrichment analysis of DEGs in classical monocytes (IMQ vs IMQ/GSK-J4 groups). Red bars indicate pathways upregulated in IMQ, and blue bars indicate pathways enriched in IMQ/GSK-J4. (E) Heatmap showing the top 50 DEGs in Tregs from the same conditions. Gene expression levels are normalized and color-coded according to relative expression intensity. (F) Volcano plot of DEGs in Tregs comparing IMQ/GSK-J4 versus IMQ groups. Volcano plots display log₂ fold change versus -log₁₀ (adjusted P). (G) GO enrichment analysis of DEGs in Tregs from IMQ vs. IMQ/GSK-J4 conditions.

To assess the impact of KDM6B inhibition on IMQ-induced inflammatory reprogramming in classical monocytes, we performed pseudobulk differential expression analysis. The heatmap representing the top 50 DEGs (Differentially Expressed Genes) in skin monocytes, selected from the union of significant DEGs (adj. p value < 0.05) across the 3 contrasts (IMQ/GSK-J4– treated vs sham–treated; IMQ-treated vs sham–treated; IMQ/GSK-J4–treated vs IMQ-treated) demonstrates that KDM6B inhibition markedly alters the transcriptional profile of classical monocytes **(Fig. 5B)**. IMQ alone induces a robust proinflammatory signature, characterized by upregulation of key mediators such as *Ifitm1, C5ar1, Saa3, Mmp8, Trem1/3, Lcn2, Lrg1, Cxcl3,* and *Cd38*. In contrast GSK-J4 significantly suppressed this hyperinflammatory program, shifting the monocyte phenotype toward a more homeostatic state **(Fig. 5B)**.

The volcano plot **(Fig. 5C)**, comparing classical monocytes from IMQ/ GSK-J4 versus IMQ treatment, highlights this transcriptional reprogramming. A total of 563 genes were detected as significantly differentially expressed (adj. P < 0.05). Among these, several proinflammatory genes were significantly downregulated (e.g., *Ifitm1, S100a8, Cxcl3, Fpr1, Lrg1, Cd38, Lcn2,* and *Acod1*), while genes associated with antigen presentation (e.g., *H2-Aa, H2-Ab1, Slamf8, Cita,* and *Mgl2*) were upregulated, indicating restoration of a regulatory state **(Fig. 5C)**. Importantly, the heatmap of adjusted p-value–filtered DEGs (n = 53) reveals that GSK-J4 treatment not only reverses IMQ-induced upregulation but also enhances expression of genes linked to antigen processing **(Fig. 5C)**.

GO analysis of classical monocytes revealed that GSK-J4 treatment significantly downregulated pathways involved in cytokine-mediated signaling (GO:0001960), TNF-α (GO:0032760) and IL-1β production (GO:0032731), canonical glycolysis (GO:0061621), regulation of monocyte differentiation (GO:0045655), complement receptor-mediated signaling pathway (GO:0002430), and neutrophil chemotaxis (GO:0030593), all of which were prominently upregulated in IMQ-treated skin **(Fig. 5D)**. In contrast, gene sets related to antigen presentation via MHC class II (GO:0002503, GO:0019886) and T cell activation (GO: 0050870) were positively enriched in the GSK-J4 group relative to IMQ **(Fig. 5D),** suggesting a reprogramming of monocytes toward a more immunoregulatory profile. These data suggest that pharmacological blockade of KDM6B shifts classical monocytes away from a hyperinflammatory phenotype toward a tolerogenic surveillance role, which may mitigate tissue damage while preserving immune function.

Further, we analyzed Treg gene-expression profiles and visualized the union of significant DEGs (adj. P < 0.05) across the three contrasts in a heatmap **(Fig. 5E)**. IMQ stimulation induced upregulation of several activation and effector markers in Tregs, including *Junb, Egr2,* and *Tnfrsf9*, suggesting a heightened activation state in response to the local inflammatory environment. Notably, genes like *Junb* and *Gzmb*, associated with Treg suppressive capacity, were also modestly upregulated in the IMQ condition, possibly reflecting a compensatory regulatory response **(Fig. 5E)**. The volcano plot highlights the top 50 significantly altered genes in skin Tregs (p-value < 0.05) **(Fig. 5F)**. Among the upregulated genes following GSK-J4 treatment were several associated with regulatory functions, including *Itgb8,* and *Il10*, while *Traf2* was significantly downregulated, suggesting an induction of anti-inflammatory programs. Conversely, key genes upregulated by GSK-J4 included proliferation-associated factors and regulatory genes such as *Mki67, Ebi3, Ccl5,* and *Socs1* **(Fig. 5F)**.

Comparison of Treg from IMQ-treated mice to sham controls showed significant enrichment of inflammatory processes, including regulation of type II interferon production (GO:0032649), positive regulation of acute inflammatory response (GO:0002675), and pathways related to CD4^+^ and CD8^+^ alpha-beta T cell differentiation (GO:0043374 and GO:0043367) **(Fig. 5G)**. Prominently, treatment with the GSK-J4 in IMQ-treated mice led to a reversal of many of these IMQ-induced inflammatory pathways. Specifically, the enrichment of type II interferon (GO:0032649) and IL-6 regulatory pathways (GO:0032715) was markedly suppressed **(Fig. 5G)**. Additionally, GSK-J4 treatment attenuated IMQ-induced enrichment of chemokine-mediated signaling (GO:0070098) and TNF response (GO:0071356), further supporting an anti-inflammatory shift in Treg function. Together, these data demonstrate that KDM6 inhibition via GSK-J4 significantly alters the transcriptional landscape of Tregs in inflamed skin, skewing them away from inflammatory phenotypes and toward a regulatory state.

### 3.6. KDM6B reduces the repressive H3K27m3 on *Il1b* and *Tnf*–mediated inflammatory cytokine promoters in monocytes

To further dissect the mechanisms underlying the pathogenic activation of monocytes in IMQ-induced psoriasis, we performed pathway enrichment analysis on differentially expressed genes (DEGs) identified in the classical monocyte populations. Further analysis showed the upregulation of gene sets associated with the regulation of TNF-α production (GO:0032760) and IL-1β production (GO:0032731) in classical monocytes from the skin of the IMQ-psoriasis mouse model **(Fig. 6A)**. Next, we investigated whether systemic increases of KDM6B contribute to BM monocyte-driven inflammation in the IMQ-psoriasis mouse model and evaluated whether GSK-J4 treatment could attenuate this heightened inflammatory response. Accordingly, on day 6 of IMQ treatment, we isolated BM monocytes for analysis. We observed increased activity and expression of KDM6B in monocytes from IMQ-treated mice compared to sham controls **(Fig. 6B)**. Additionally, levels of intracellular α-KG [33] were elevated in BM monocytes from IMQ-treated mice **(Fig. 6B)**. These increases were associated with heightened expression of *Il1b* and *TNF*, which are crucial pro-inflammatory cytokines regulated by NF-κB, at both the transcriptional and protein levels, which are known contributors to psoriatic skin inflammation **(Fig. 6C)**. Notably, GSK-J4 treatment significantly reduced KDM6B activity, lowered *Kdm6b* expression, partially normalized intracellular α-KG, and consequently the expression of IL-1β and TNF-α, demonstrating its effectiveness in mitigating the inflammatory response **(Figs. 6A-6C)**.

**Fig. 6.**
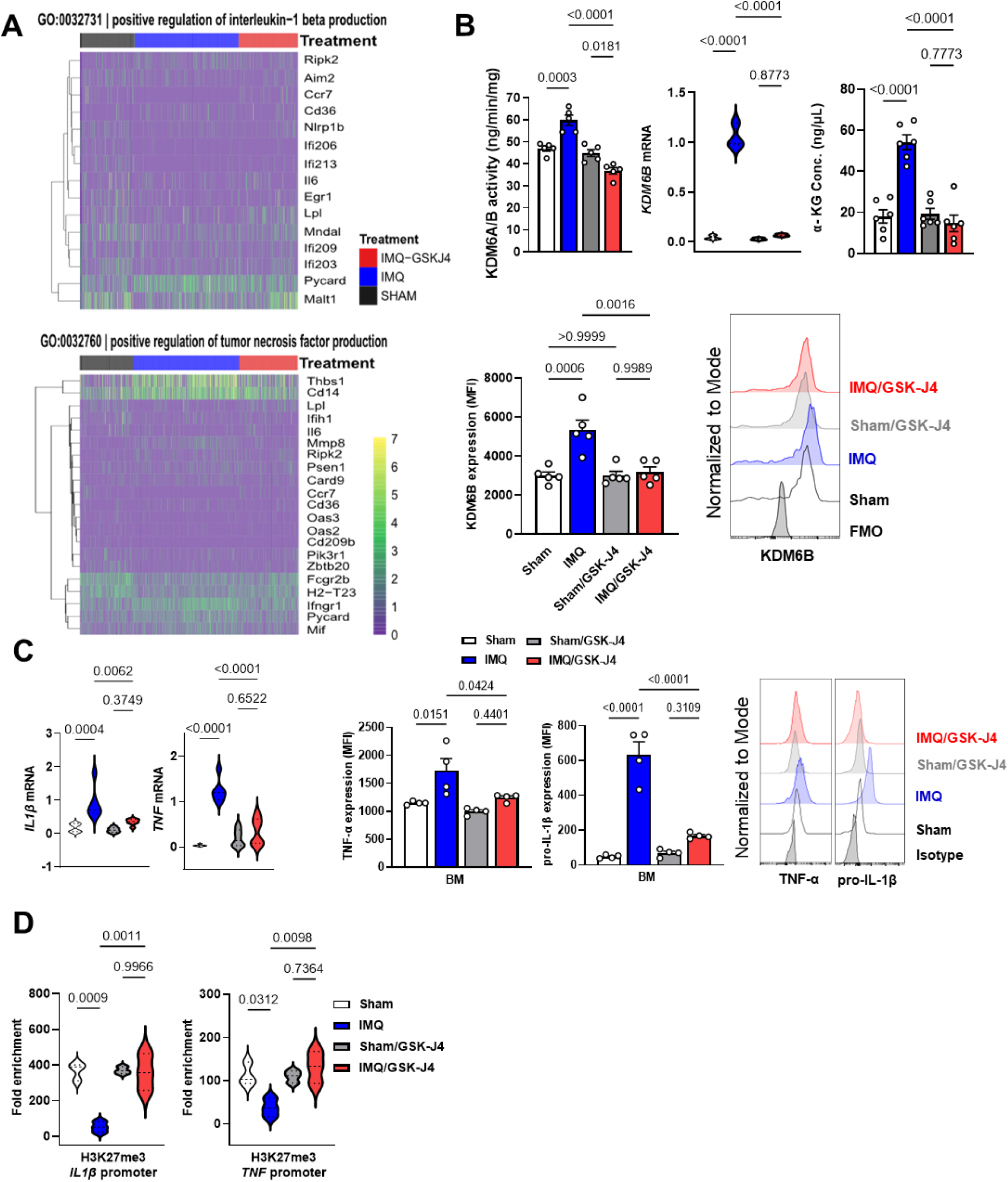
KDM6B is increased in IMQ-psoriasis monocytes and reduces the repressive H3K27me3 on *Il1b* and *TNF* cytokine gene promoters. (A) Heatmaps of differentially expressed genes in classical monocytes isolated from skin of sham-, IMQ-, and IMQ+GSK-J4–treated mice. Shown are representative inflammatory genes that involve TNF-α production (GO:0032760) and IL-1β production (GO:0032731) with color scales indicating relative expression levels. The color scale indicates standardized expression levels (z-scores), with yellow representing higher expression and violet representing lower expression relative to the mean. (B) KDM6B enzymatic activities and KDM6B gene expression were measured by quantitative PCR, flow cytometry, and ELISA. α-KG levels in BM monocytes from IMQ-treated mice with and without GSK-J4 compared to sham-treated mice. (C) *Il1b* and *Tnf* gene expressions were measured by quantitative PCR in BM monocyte from IMQ-treated mice with and without GSK-J4 compared to sham-treated mice. (D) ChIP–qPCR analysis of H3K27me3 at *Il1b* and *Tnf* promoters in BM monocytes from sham-, IMQ-, GSK-J4-, and IMQ/GSK-J4-treated mice; isotype-matched IgG controls run in parallel; signals normalized to input. Data are representative of three independent experiments (n = 4-6) and are presented as mean ± SEM. Statistical significance was determined using one-way ANOVA with the Newman–Keuls multiple comparison test.

Mechanistically, KDM6B functions through H3K27me3 demethylation at proinflammatory gene promoters [14,41]. Elevated KDM6B activity reduces H3K27me3, thereby enhancing the accessibility of NF-κB to promoter regions and activating gene transcription [34,40]. To directly examine this, chromatin immunoprecipitation qPCR (ChIP-qPCR) assays were performed on BM monocytes from IMQ-treated mice compared to sham-treated controls. ChIP-qPCR analysis using primers targeting NF-κB binding sites on *Il1b* and *Tnf* gene promoters demonstrated reduced H3K27me3 occupancy at these sites in BM monocytes from IMQ-treated mice **(Fig. 6D)**. Importantly, GSK-J4 treatment significantly reversed these changes, decreasing KDM6B activity, KDM6B expression, IL-1β and TNF-α gene expression, and restoring H3K27me3 levels at NF-κB binding sites of *Il1b* and *Tnf* gene promoter regions. While monocytes are likely the primary targets in this context, we cannot exclude potential effects of GSK-J4 on other immune cell subsets. Collectively, these results support a causative role for KDM6B in driving IL-1β and TNF-α–mediated inflammation in monocytes during IMQ-psoriasis inflammation.

### 3.7. GSK-J4 reverses glycolytic metabolic remodeling in classical monocytes in the IMQ-psoriasis mouse model

Recent studies indicate that reprogramming of cellular metabolism and epigenetic modifications can regulate monocyte phenotype and function [12,42]; however, little is known about the metabolic programs that characterize monocytes in IMQ-psoriasis. GO analysis identified canonical glycolysis among the most significantly enriched pathways in IMQ-treated versus sham-treated classical monocytes (**Fig. 7A)**. Specifically, 10 out of 18 canonical glycolysis-related genes (GO:0061621), including *Pkm, Eno1, Aldoa, Gapdh, Pgk1, Pgam1, Tpi1, Gpi1, Pfkp, and Pfk1,* were upregulated in classical skin monocytes from IMQ-treated mice compared to sham controls **(Fig. 7A)**. Additionally, the gene expression of these glycolytic enzymes was significantly increased in BM monocytes of IMQ-treated mice compared to sham-treated controls **(Fig. 7B)**. Mice treated with IMQ in combination with GSK-J4 displayed a significant reduction in the expression of canonical glycolytic enzyme genes, including *Pgam1*, *Pgk1*, *Aldoa*, *Pkm2*, and *Eno1*, in BM monocytes **(Fig. 7B)**.

**Fig. 7.**
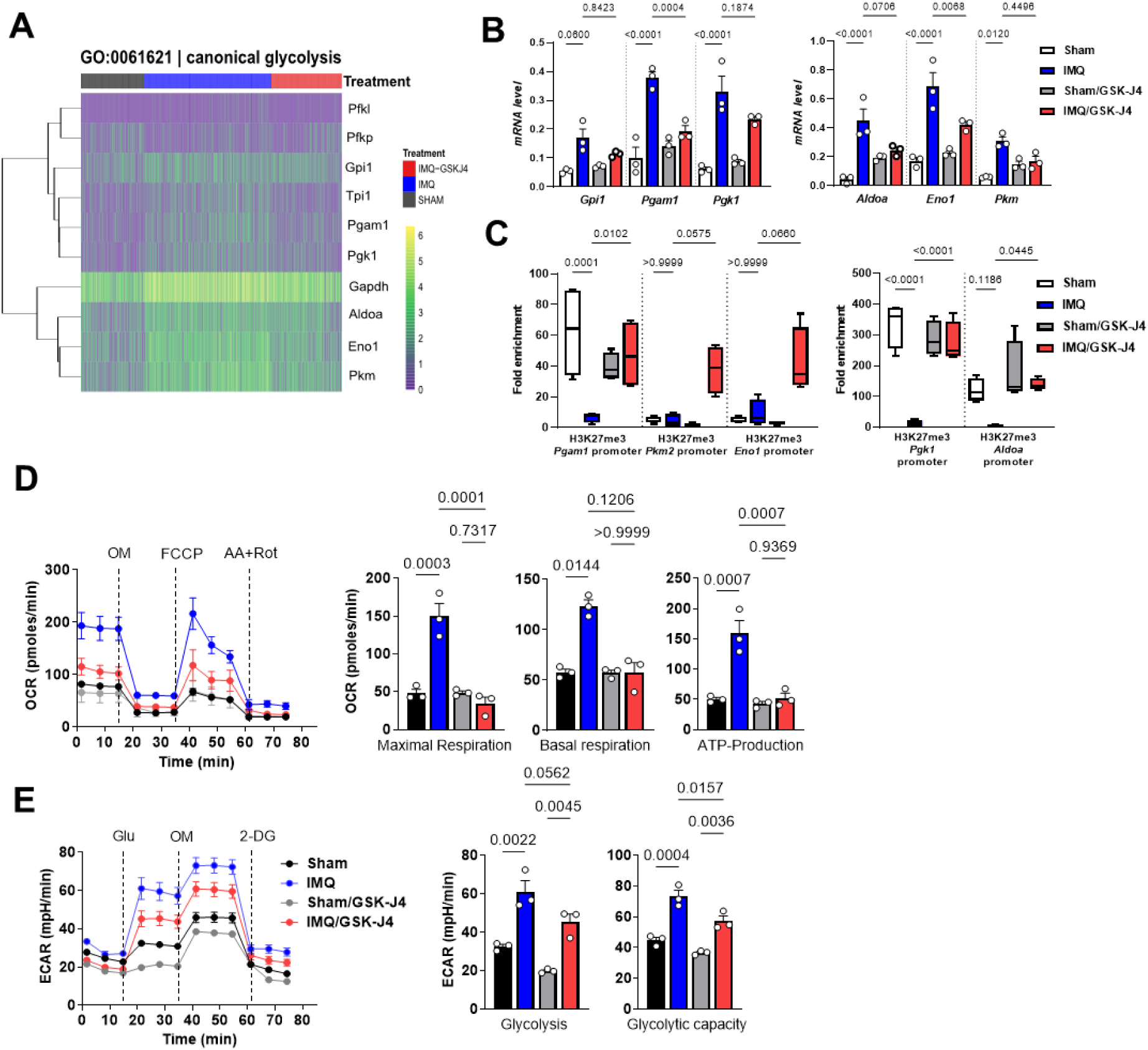
Metabolic remodeling of inflammatory monocytes in IMQ-psoriasis mouse model. (A) Heatmap of gene expression of canonical glycolysis enzymes in classical skin monocytes from IMQ-treated compared to sham- and IMQ/GSK-J4-treated mice. The color scale indicates standardized expression levels (z-scores), with yellow representing higher expression and violet representing lower expression relative to the mean. (B) Gene expression levels of key glycolytic enzymes in BM-derived monocytes, assessed by RT-qPCR. (C) ChIP-qPCR analysis of H3K27me3 enrichment at the promoters of glycolytic genes in BM monocytes from IMQ-treated mice with and without GSK-J4 compared to sham-treated mice. (D) Seahorse analysis of cellular OCR, basal respiration, maximum respiration, and spare respiratory capacity. OM: oligomycin; FCCP: carbonyl cyanide-4-(trifluoromethoxy)phenylhydrazone (uncoupler); AA+Rot: antimycin A + rotenone. (E) Seahorse analysis of ECAR, glycolysis, glycolytic capacity, and glycolytic reserve, in BM monocytes from IMQ-treated mice with and without GSK-J4 compared to sham-treated mice with sequential injections of glucose (Glu), oligomycin (OM), and 2-deoxy-D-glucose (2-DG). Data are representative of three independent experiments (n = 3–5) and are presented as mean ± SEM. Statistical significance was determined using one-way ANOVA with the Newman–Keuls multiple comparison test.

To further investigate whether KDM6B regulates these metabolic genes via H3K27me3 demethylation at their promoter regions, we performed ChIP-qPCR analysis for H3K27me3 enrichment at selected glycolytic gene promoters. In BM monocytes from IMQ-treated mice, we observed a marked reduction in repressive H3K27me3 marks at the promoters of *Pgam1*, *Pgk1*, and *Aldoa*, indicating active KDM6B-mediated demethylation **(Fig. 7C)**. Importantly, GSK-J4 treatment restored H3K27me3 enrichment at the promoters of *Pgam1*, *Pgk1*, and *Aldoa*, and partially at *Pkm2* and *Eno1* **(Fig. 7C)**, suggesting that pharmacological inhibition of KDM6B reverses epigenetic activation of glycolytic programs in classical monocytes. Collectively, these results support a causative role for KDM6B in promoting glycolytic reprogramming of BM monocytes during IMQ-induced psoriasis, through locus-specific H3K27me3 demethylation at key metabolic gene promoters.

Metabolic reprogramming from oxidative phosphorylation (OXPHOS) toward glycolysis is essential for monocytes to effectively initiate and sustain pro-inflammatory responses [12,43]. To systematically characterize the metabolic activity of monocytes from IMQ-treated mice, we measured the oxygen consumption rate (OCR) and extracellular acidification rate (ECAR) in BM-derived monocytes, indicative of OXPHOS and glycolytic activities, respectively. Consistent with observations in monocytes from psoriasis patients [17] and the enrichment of canonical glycolysis-related genes **(Fig. 7A)**, IMQ-treated monocytes displayed elevated basal and maximal OCR as well as spare respiratory capacity, reflecting enhanced mitochondrial respiration. Treatment with GSK-J4 significantly reduced OCR parameters, suggesting lower cellular energy demands associated with diminished inflammatory activation **(Fig. 7D)**. Furthermore, IMQ-treated BM monocytes exhibited increased ECAR, indicative of heightened glycolytic activity and glycolytic capacity, both of which were significantly attenuated by GSK-J4 treatment **(Fig. 7E)**. Collectively, these results indicate that monocytes migrating from BM to inflamed skin undergo substantial metabolic remodeling to fulfill increased bioenergetic demands during inflammation. These data support a model in which KDM6B activity sustains both glycolytic and mitochondrial programs during inflammatory activation, and where pharmacologic inhibition rebalances monocyte metabolism toward a less inflammatory state.

## 4. Discussion

Our study demonstrates that the histone demethylase KDM6B functions as an epigenetic-metabolic checkpoint in classical monocytes that promotes psoriasis-induced inflammation. Peripheral depletion of classical Ly6C^hi^ monocytes or pharmacological blockade of KDM6B with GSK-J4 reduced both skin and systemic inflammation in the IMQ-psoriasis mouse model, even when treatment began after disease onset. Upregulation of KDM6B removes the repressive H3K27me3 marks from NF-κB-responsive *Il1b/Tnf* promoters and key glycolytic genes (*Pgam1, Pgk1,* and *Aldoa*), linking chromatin opening to enhanced glycolytic flux and elevated inflammatory cytokine production. Conversely, the KDM6B inhibitor GSK-J4 restores H3K27me3, suppresses both the metabolic and inflammatory pathways, and normalizes monocyte metabolism. Previous work suggested that psoriasis is a disease primarily driven by keratinocyte-T cell crosstalk [1,4,44]. Our data now shows a crucial role of BM-derived monocytes in IMQ-induced inflammation, as they infiltrate lesional skin in large numbers, adopt a transcriptional program dominated by NF-κB, and secrete IL-1β and TNF-α in a KDM6B-dependent manner. The rise in α-ketoglutarate, an essential cofactor for KDM6B demethylases, indicates that inflammatory metabolism actively preserves the permissive chromatin state. This, in turn, generates a feed-forward loop in which metabolism reinforces the underlying chromatin landscape that drives inflammation. Blocking KDM6B breaks this loop, limits neutrophil recruitment, and is accompanied by an increase in FOXP3^+^ regulatory T cell numbers, implying that monocyte reprogramming has downstream effects on adaptive immunity.

Recent studies show that KDM6B drives maladaptive myeloid reprogramming across multiple inflammatory diseases [18,35,45,46]. *Davis et al*. identified KDM6B as a central regulator of monocytes/macrophage-driven inflammation in the development of abdominal aortic aneurysms (AAAs) [14]. Both human AAA specimens and mouse models showed markedly increased KDM6B expression in monocytes and macrophages infiltrating the aortic wall [14]. In diabetic wounds, excessive KDM6B activity impairs reparative monocyte and macrophage function through aberrant activation of the cGAS–STING pathway [41], whereas in glioblastoma, myeloid-specific Kdm6b deletion or pharmacological inhibition with GSK-J4 restores antigen presentation, enhances type I interferon signaling, and sensitizes tumors to anti-PD-1 therapy [13]. Mechanistically, KDM6B promotes inflammation by removing the repressive H3K27me3 mark on promoters of NF-κB target genes such as *Il1b, Il12,* and *Tnf*, thereby allowing their transcription. Upstream, type I interferon (IFN-β) induces KDM6B expression via the JAK1/STAT1 pathway [14]. Audu et al. further showed that KDM6B acts as an epigenetic ‘gatekeeper’, deciding whether wound-resident macrophages resolve or prolong inflammation. During normal repair, a transient IFN-β increase activates JAK1/3-STAT3 signaling early after injury, producing a short-term increase in KDM6B. In obesity-associated diabetic wounds, however, persistent IL-6 maintains the same pathway, generating a delayed but sustained KDM6B induction that removes H3K27me3 from *Il1b, Tnf,* and the cGAS-STING gene *Tmem173* and amplifies NF-κB signaling [41]. In systemic SLE monocytes, chronic type I IFN exposure elevates intracellular α-ketoglutarate, boosts KDM6B activity, reduces H3K27me3 at interferon-stimulated gene promoters and sustains an inflammatory “trained-immunity” state [18]. Our data extend this paradigm to the skin and place psoriasis within the broader spectrum of trained-immunity disorders in which chromatin and metabolism are coopted to sustain innate immune memory in BM. Consistent with findings in SLE [18], GSK-J4 treatment in the IMQ model of psoriasis restores repressive H3K27me3 and dampens pathogenic transcriptional programs, emphasizing the translational potential of targeting KDM6B in autoinflammatory disease.

Despite its efficacy, GSK-J4 lacks absolute selectivity and can inhibit non-histone proteins and the related demethylase KDM6A at higher concentrations [16,35]. Moreover, the specific contribution of monocytes to psoriasis-induced systemic inflammation remains to be defined. Systemic KDM6B inhibition is likely to impact multiple immune and non-immune lineages, including Th17 cells, dendritic cells, and keratinocytes. Since earlier studies showed that KDM6B inhibition prevents Th17 differentiation, a cell-type-specific Kdm6b-deficient mouse would be invaluable for examining these lineage-specific effects [39,47]. A further limitation is that the IMQ-psoriasis mouse model reflects only an acute inflammatory response, highlighting the need to investigate chronic psoriasis models as well.

Despite the success of IL-17 and IL-23 biologic therapies, up to one-third of patients remain non-responders or eventually relapse [48]. Resetting the innate-immune threshold upstream of cytokine production through KDM6B inhibition could potentiate these agents, extend dosing intervals, and restore responsiveness in treatment-refractory patients. Emerging modalities such as locus-selective protein degraders and allosteric inhibitors promise greater specificity than GSK-J4 while retaining comparable anti-inflammatory efficacy.

Collectively, our findings position KDM6B as a central regulator linking chromatin rewiring to metabolic reprogramming in proinflammatory monocytes and show that pharmacological blockade of this demethylase restores both innate and adaptive immune responses in established psoriatic disease. Therapeutically targeting the KDM6B–H3K27me3 axis thus offers a tractable approach to disrupt the self-sustaining loop of epigenetic and metabolic priming that drives psoriasis and other myeloid-mediated autoinflammatory disorders.

## CRediT authorship contribution statement

**Aman Damara:** Data curation, Methodology, Formal analysis, Writing – review & editing. **Najla Abassi:** Data curation, Formal analysis, Writing – review & editing. **Delia Mihoc:** Data curation, Writing – review & editing. **Mahsa Nastaranpour:** Data curation. **Pauline Kraft:** Data curation. **Tina Sarkar:** Data curation. **Carsten Deppermann:** Writing – review & editing. **Johannes U Mayer:** Writing – review & editing. **Stephan Grabbe:** Writing – review & editing. **Michael Delacher:** Writing – review & editing, Resources, Supervision. **Federico Marini:** Writing – review & editing, Resources, Supervision. **Fatemeh Shahneh:** Writing – review & editing, Writing – original draft, Resources, Supervision, Project administration, Funding acquisition, Formal analysis, Data curation, Conceptualization.

## Acknowledgments

We thank the FZI NGS and Flow Cytometry Core Facility, as well as the Animal Facility of the University Medical Center Mainz, for their excellent technical support.

## Data availability

We provide the annotated single-cell RNA-seq datasets via an interactive application, built with the iSEE package and available at http://shiny.imbei.uni-mainz.de:3838/iSEE_KDM6B_inhibition.

