## Supplemental Fig 1 and 2 for "KDM6B inhibition modulates monocyte activation and alleviates IMQ-psoriasis skin inflammation"

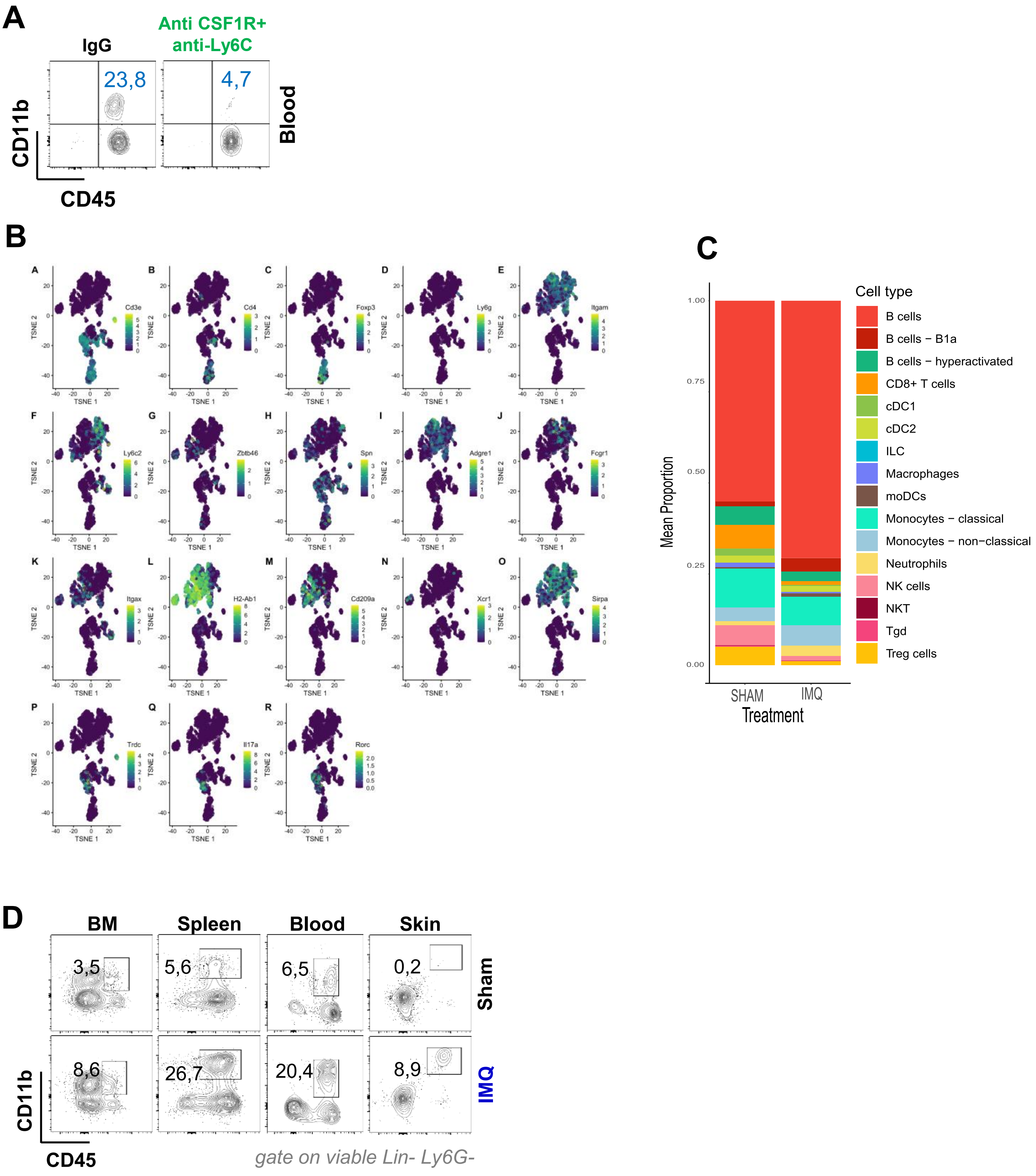


Fig. S1. (A) Representative flow cytometry plots of peripheral blood one day after a single dose: isotype IgG control and anti-CSF1R + anti-Ly6C–treated. (B) t-SNE plots of marker genes used to identify each immune-cell cluster. (C) Stacked bar plot showing the relative abundance of immune cell subsets in the spleen of sham- versus IMQ-treated mice. (D) Flow-cytometric analysis of $CD11b^{+}Ly6G^{-}$ monocytes in BM, spleen, blood, and skin.

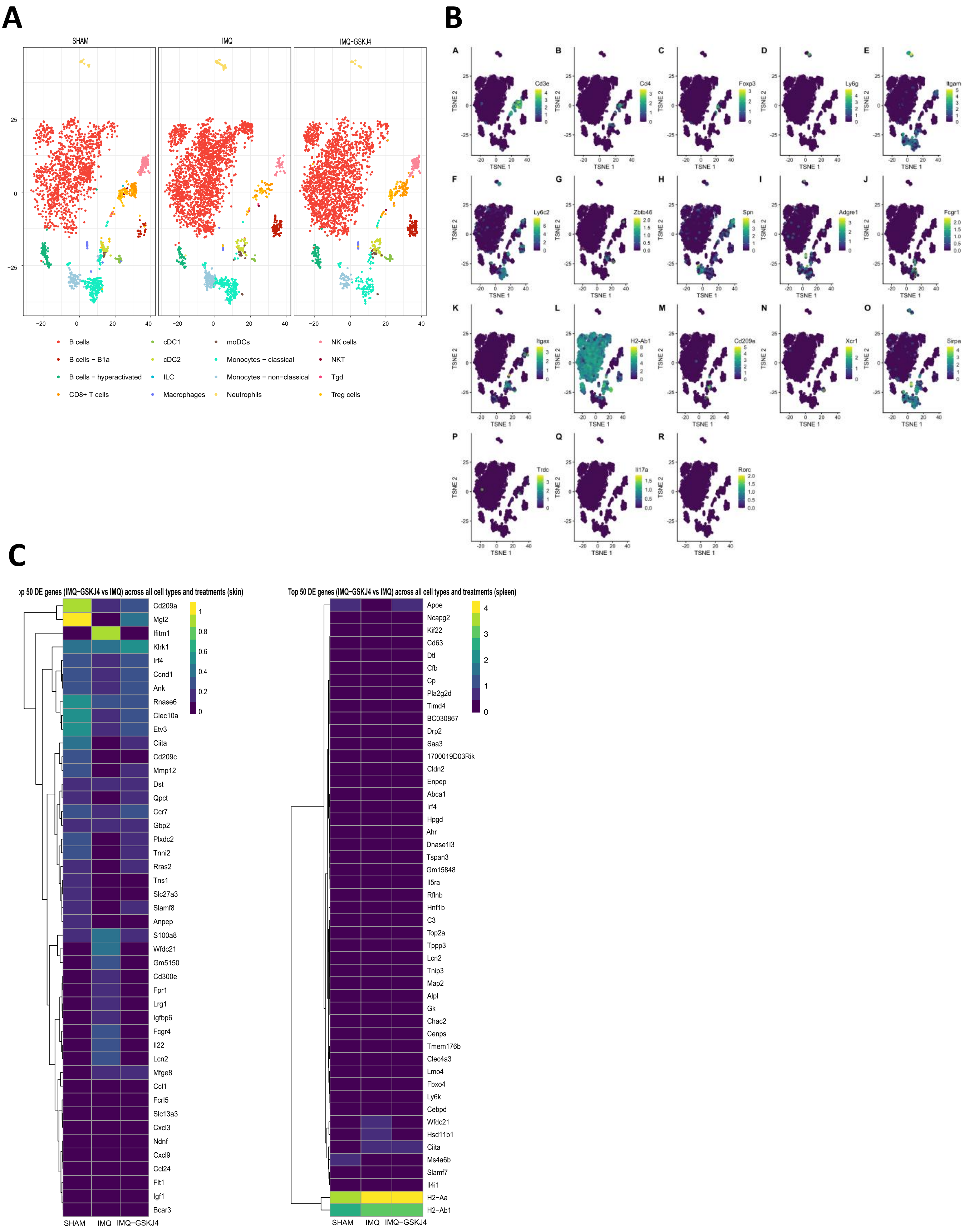


Fig. S2. (A) t-SNE map of CD45$^{+}$ immune cells isolated from the spleen from sham-treated, IMQ-treated, and IMQ/GSK-J4-treated groups. (B) t-SNE feature plots of canonical marker genes used to identify each immune-cell cluster. (C) Heatmap of pseudobulk expression of the top 50 DEGs across all cell types in skin and spleen from sham, IMQ, and IMQ/GSK-J4 groups.
